# Cleaning the Dead: Optimized decontamination enhances palaeoproteomic analyses of Pleistocene skeletal material

**DOI:** 10.1101/2024.06.13.598810

**Authors:** Zandra Fagernäs, Viridiana Villa Islas, Gaudry Troché, Jan-Pieter Buylaert, Tura Khujageldiev, Redzhep Kurbanov, Jesper V. Olsen, Mikkel Winther Pedersen, Frido Welker

## Abstract

The study of ancient proteins preserved in a range of archaeological, cultural heritage, and palaeontological materials is increasingly contributing to our understanding of human evolution and archaeological research questions. Many of the specimens studied have been excavated and stored for a significant duration prior to their proteomic analysis. Human handling and storage environments therefore provide ample opportunities for protein contamination onto and into specimens of interest to palaeoproteomic studies. As such, modern protein contamination limits access to endogenous proteomes. Here, we compare five approaches of bone protein decontamination applied to a Pleistocene *Equus* sp. bone fragment contaminated with a modern dog salivary proteome. We find that all tested methods reduce the protein contamination, but with different efficiencies. We find that a brief bleach wash is the most effective approach in removing modern protein contamination, and that no additional damage is caused to the endogenous proteome by this treatment. Next, we apply this approach to a hominin tooth found at Khudji, a Late Pleistocene archaeological site in Tajikistan. We demonstrate that a brief bleach wash removes almost all human skin protein contamination while retaining the endogenous hominin dentine proteome. Subsequent phylogenetic analysis of the Khudji dentine proteome allowed determination that the specimen is likely not a Denisovan, but still leaves ambiguity between an assignment to either modern humans or Neanderthals.

## 1. Introduction

Palaeoproteomics is a fast-growing field within the archaeological sciences, heritage studies, and palaeoanthropology, with applications such as phylogenetic studies of extinct taxa (Cappellini et al., 2019; Welker et al., 2019), human evolution (Chen et al., 2019; Welker et al., 2020), human subsistence behavior (Le Meillour et al., 2020; Sinet-Mathiot et al., 2019), and health and disease (Fotakis et al., 2020; Warinner et al., 2014), among others. As proteins have been found to preserve in a diverse range of archaeological substrates, and can preserve for millions of years (Demarchi et al., 2022; Madupe et al., 2023), new areas of archaeological studies continue to be opened up as new data types are generated. However, the study of ancient proteins is not straightforward, as the proteome is significantly altered through time. Over time proteins are fragmented into increasingly shorter peptides, resulting in some proteins being completely lost (Welker, 2018). Simultaneously, the fragmented but surviving protein sequences contain amino acids with increasing levels of damage, in different forms (Hendy, Welker, et al., 2018; Warinner et al., 2022). Additionally, contaminating proteins are added to the ancient proteome, deriving from sources such as the burial environment, excavation, storage and subsequent handling in curatorial facilities, and potentially in laboratory and mass spectrometry environments as well (Hendy, Welker, et al., 2018). If not removed or identified, these contaminating proteins risk negatively impacting downstream analyses by i) masking endogenous proteins due to their abundance and more intact state, and ii) facilitating the unwanted incorporation of modern protein sequence variation into reconstructed ancient protein sequences.

Significant efforts have been dedicated to the introduction of extraction and injection blanks during palaeoproteomic laboratory workflows. Extraction blanks within the laboratory environment serve to control for the introduction of any protein residues during the extraction process, while injection blanks on protein mass spectrometry equipment are meant to control for the introduction or carryover of peptides within the mass spectrometer (Demarchi et al., 2022; Hendy, Welker, et al., 2018). Although these are necessary, neither of these controls can, however, provide assessment of the endogenous or contaminant origin of proteins and peptides deposited onto a palaeoproteomics study object prior to protein extraction.

Additional research on the separation of contaminants and endogenous peptides and proteins has focused on the bioinformatic analysis of diagenetic modifications at the amino acid or peptide level, under the assumption that some of these will show different quantitative or qualitative properties between peptides originating from contaminating proteins compared to endogenous ones. Examples of these include the quantification of the extent of asparagine (N) and glutamine (Q) deamidation, quantification of the extent of various oxidative amino acid modifications, the calculation of peptide length distributions, and analysis of peptide terminus states (Cappellini et al., 2019; Chen et al., 2019; Mackie et al., 2018; Orlando et al., 2013; Ramsøe et al., 2020). Often, metrics derived from these measurements are averaged across the full proteomes identified, while in some cases sufficient amounts of data is available to perform these assessments per protein group (Chen et al., 2019; Ramsøe et al., 2021; Welker et al., 2020). These bioinformatics approaches are useful and necessary when studying ancient skeletal proteomes, but are not without problems. They generally require a sufficiently large number of PSMs per protein group, which may not be available for most protein groups. They rely on some quantifying metric, which at times might be difficult to establish confidently. Furthermore, extraction methods themselves might introduce modifications to the extracted contaminating proteins, mimicking the effect of diagenesis. Finally, none of these approaches resolve the effect of abundant modern proteins masking the identification of degraded and low-abundance proteins from ancient skeletal specimens.

In parallel to the development of bioinformatics approaches to characterise the diagenetic modification of proteins in palaeoproteomics contexts, a number of studies therefore mention approaches to decontaminate an archaeological sample prior to protein extraction. Such claimed methods include mechanical surface removal (Kontopoulos et al., 2020; Sawafuji et al., 2017; Wasinger et al., 2019), washing with bleach (sodium hypochlorite) (Froment et al., 2020; Sakalauskaite et al., 2020; Trolle Jensen et al., 2020), washing with ethylenediaminetetraacetic acid (EDTA) (Fagernäs et al., 2020; Hendy, Colonese, et al., 2018; Sawafuji et al., 2017), washing with hydrochloric acid (HCl) (Gasparini et al., 2022; Palmer et al., 2021; Wasinger et al., 2019), washing with water (Gasparini et al., 2022; Spengler et al., 2022), and UV irradiation (Fagernäs et al., 2020; Froment et al., 2020). Although a wide range of published protocols therefore include some kind of decontamination step, there is no consensus on which decontamination method should be used. In fact, there are no comparisons of the efficiency of the above-mentioned approaches in removing protein contamination, nor has their impact on the endogenous proteome been studied.

Here, we compare five decontamination methods for the palaeoproteomic analysis of an artificially contaminated Pleistocene *Equus* sp. bone specimen; washing with bleach, HCl, EDTA or water, and UV irradiation. These methods were chosen based on their frequent application in palaeoproteomic studies, regardless of proven or theoretical efficiency. After determining that a mild bleach treatment is, of the compared approaches, the most suited for removing modern protein contamination, we then apply this approach to dentine from a Pleistocene hominin tooth from Khudji, Tajikistan, which was found to be heavily contaminated with proteins deriving from human skin. Bleach treatment removed nearly all contaminating proteins, without damaging the endogenous hominin proteins present.

## 2. Material and methods

### 2.1. Samples

A Pleistocene faunal bone fragment from the Dutch North Sea shore was used to compare different decontamination methods. Previous palaeoproteomic analysis had shown that this fragment stems from an *Equus* sp. A sample was removed from the bone specimen by drilling after manual contamination by contact with saliva and skin of a dog (Figure S1). Specifically, saliva was manually transferred from the dog’s mouth using a nitrile glove, and smeared over the bone, whereafter the bone was rubbed on the dog’s back. The contamination was not performed in a clean laboratory environment but instead in an office environment, potentially introducing both human and dog proteins. This is not unlike experimental data that suggests the consistent recovery of skin, hair, and salivary proteins from archaeological skeletal specimens, and which are generally regarded as representing modern protein contamination. The dog had not eaten recently prior to the contamination and had no known oral disease.

The most promising decontamination method was applied to a hominin tooth from the archaeological site of Khudji, Tajikistan (Figure 6A; Supplementary Information A), which showed a large amount of human skin contamination during initial analyses. This is a deciduous incisor, excavated in 1997, from a layer radiocarbon dated to approximately 40,000 years before present. The specimen has an unknown taxonomic identity within the genus *Homo* based on morphological analyses (Trinkaus et al., 2000).

### 2.2. Laboratory methods

All laboratory work was conducted in dedicated facilities for the processing of ancient biomolecules at the Globe Institute, University of Copenhagen, Denmark. Extraction blanks were carried alongside samples to monitor laboratory contamination.

***Equus* sp. sample:** The part of the contaminated bone piece with highest visible amount of contamination (Figure S1) was powdered to a homogenous powder using a mortar and pestle, and divided among the different decontamination approaches (Figure 1A). Additionally, a sample was taken from the original uncontaminated bone fragment, and powdered with a mortar and pestle (Supplementary Data 1).

**Figure 1.**
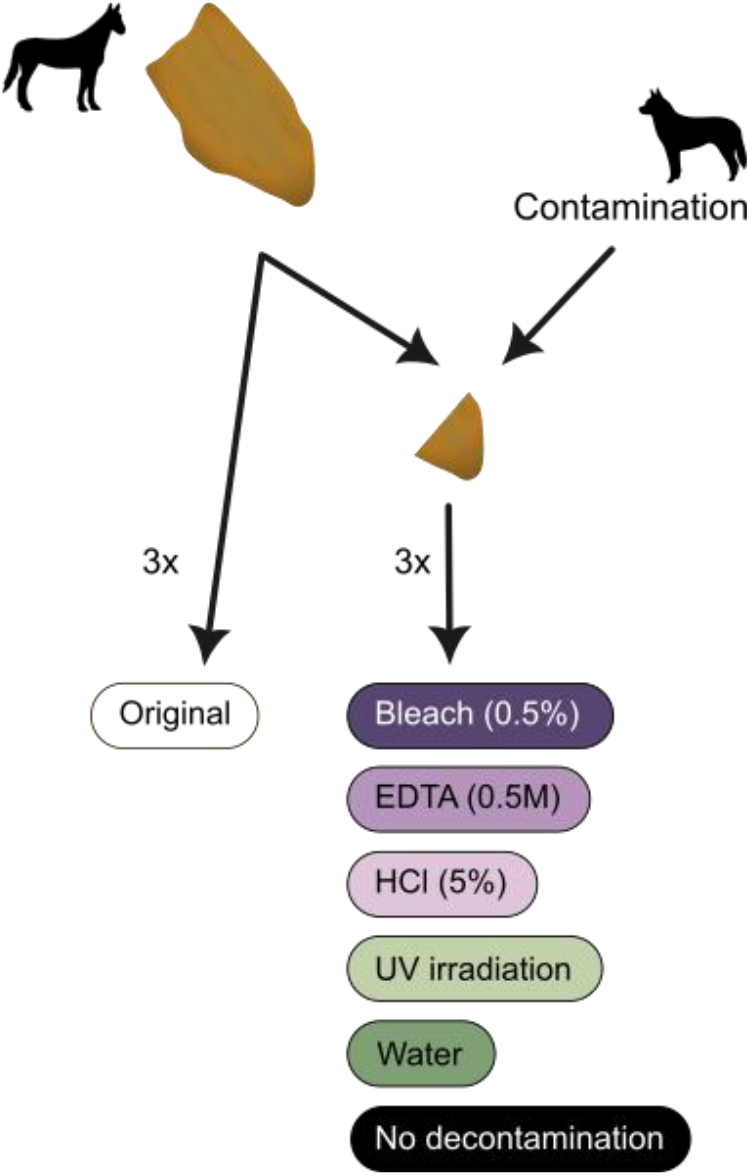
Schematic of the laboratory workflow.

Five different decontamination methods were tested: Washing with bleach (sodium hypochlorite), HCl (hydrochloric acid), EDTA (ethylenediaminetetraacetic acid), molecular grade water, and UV irradiation (Figure 1; Table 1). These methods were selected as they are commonly used for decontamination in palaeoproteomics literature. They do not represent a complete list of all possible decontamination methods, nor were they selected for their likely effectiveness. Mechanical surface removal, although commonly utilized, was not included as it requires destruction of a higher amount of sample, which is not desirable for archaeological materials. For all the methods including washes, 1 ml of the reagent was added to the sample, vortexed for 5 s, and centrifuged for 1 min at 13 krpm, whereafter the reagent was removed. After the washes with bleach and HCl, the bone pellet was washed twice with 0.5 ml molecular grade water, in order to avoid the reagents interfering with downstream analyses. UV irradiation was conducted in a crosslinker (UVP Crosslinker, Analytik Jena) with the bone powder in an open 2 ml Eppendorf tube. After 30 s of irradiation, the tube was shaken, and the irradiation continued for an additional 30 s. Additionally, a contaminated sample was included with no decontamination performed, as well as a non-contaminated (“original”) sample. All methods outlined above were conducted in triplicate, in order to account for between-extract variation. Proteins were extracted from the bone powder using a standard palaeoproteomic protocol (Lanigan et al., 2020) and peptides were cleaned on in-house made StageTips (Supplementary Information A).

**Table 1.**
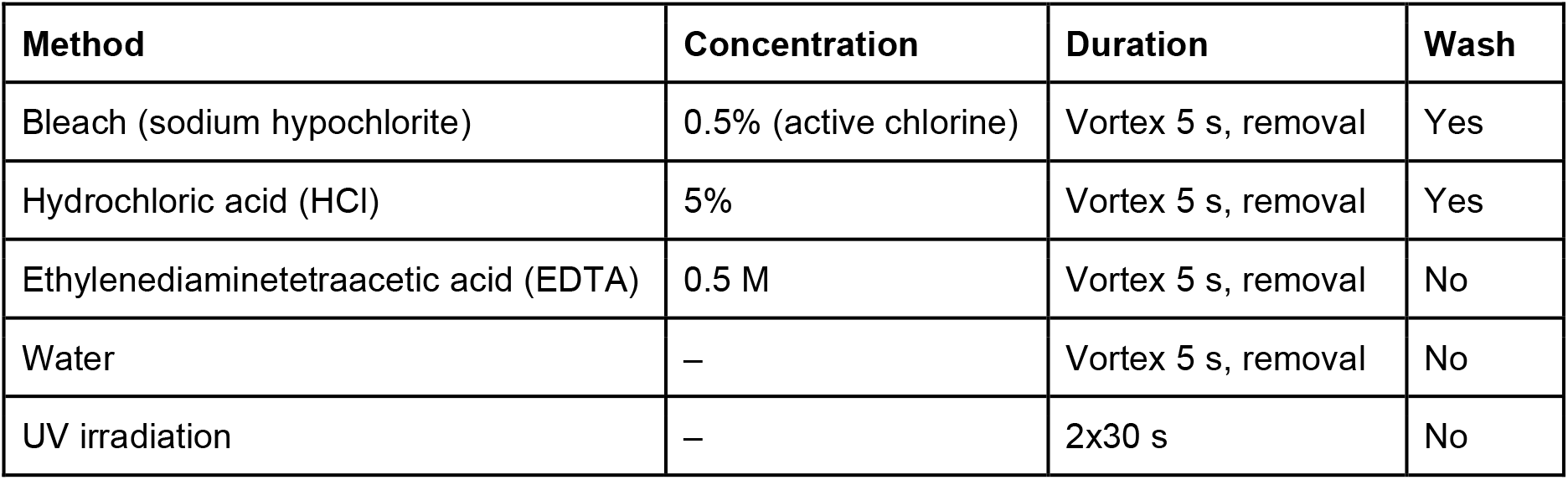
Decontamination methods included in the current study. Details shown include the concentration of the used reagent (when applicable), the duration of decontamination, and whether a wash with molecular grade water was conducted afterwards or not.

| Method | Concentration | Duration | Wash |
| --- | --- | --- | --- |
| Bleach (sodium hypochlorite) | 0.5% (active chlorine) | Vortex 5 s, removal | Yes |
| Hydrochloric acid (HCl) | 5% | Vortex 5 s, removal | Yes |
| Ethylenediaminetetraacetic acid (EDTA) | 0.5 M | Vortex 5 s, removal | No |
| Water | – | Vortex 5 s, removal | No |
| UV irradiation | – | 2x30 s | No |

***Homo* sp. sample:** A small fragment was removed from the root of the Khudji hominin incisor through drilling, and subsampled for different extractions (Supplementary Information A). Proteins were extracted using a protocol adapted for highly degraded samples (Jensen et al., 2023). For the second subsample, a bleach decontamination step was added prior to demineralization as described above.

### 2.3. Mass spectrometry

For all samples, LC-MS/MS was conducted at the Centre for Protein Research (University of Copenhagen) (see details in Supplementary Information A). An EASYnLC 1200 system was utilised for liquid chromatography (Thermo Fisher Scientific, Waltham, MA, USA) and mass spectrometric analysis was conducted using an Exploris 480 (Thermo Fisher Scientific, Waltham, MA, USA).

### 2.4. Data analysis

The *Equus* sp. raw data was analyzed using MaxQuant v.2.1.3.0 (Cox & Mann, 2008) with the dog (*Canis lupus familiaris*) and horse (*Equus caballus*) reference proteomes, as well as the internal MaxQuant contaminant database (see details in Supplementary Information A). A semi-specific search was conducted, with Oxidation (M), Deamidation (NQ), Gln/Glu->pyro-Glu, Carbamidomethylation and Hydroxyproline as variable modifications, and adding Oxidation (WH) and Dioxidation (WH) when investigating UV-damage. The Homo sp. data was analyzed using the human reference proteome with a semi-specific search and Oxidation (M), Deamidation (NQ), Gln/Glu->pyro-Glu and Hydroxyproline as variable modifications.

The *Homo* sp. proteome was reconstructed for proteins with more than 5 peptides based on a PEAKS v.7.0 (Zhang et al., 2012) search against a database of the modern human proteome with added archaic variation (see details in Supplementary Information A). A majority consensus was called for each amino acid position, and any identified SAPs were manually verified. Thereafter, a consensus was created from the two reconstructions, leading to three reconstructed proteomes in total: “Original”, “Bleach”, and “Consensus” (Table S1). Phylogenetic analysis was conducted using both a maximum likelihood method through RAxML (Stamatakis, 2006) and a Bayesian approach using MrBayes (Ronquist et al., 2012)

## 3. Results

### 3.1. Decontamination method comparison

Although previous palaeoproteomic studies have mentioned that modern protein contamination is occurring and is being identified, there is no evidence of the impact contamination has on an endogenous skeletal proteome. We therefore analyzed a Pleistocene *Equus* sp. bone that had been contaminated by modern dog saliva and skin (Figure 1). This combination was chosen to have protein contamination that is easily recognizable both taxonomically and in terms of proteome composition. Additionally, the salivary proteome is a complex and comparatively large proteome (Torres et al., 2018) that absorbs into the bone specimen, thereby mimicking a realistic contamination scenario. Human proteins might also be present due to prior handling of the specimen in curatorial contexts, as well as having been added during the dog protein contamination event.

We first analyzed the extracted proteomes from the contaminated, non-decontaminated samples (n=3) together with the original, uncontaminated samples (n=3) against the *Equus caballus* (horse) proteome (including the MaxQuant contaminant database). Our results show that the horse proteome identified is different when comparing these two conditions, with the conditions clearly separated in PCA space (Figure 2A). The original samples have 25.0±3.0 (mean ± standard deviation) Equus proteins and 1,439.3±97.8 PSMs identified, whereas the contaminated samples have 64.0±5.3 proteins and 1,019.3±74.7 PSMs. The contaminated proteome that is reconstructed thereby has a larger number of proteins, but with fewer PSMs, and includes salivary proteins that are assigned as horse since a dog database is not included.

**Figure 2.**
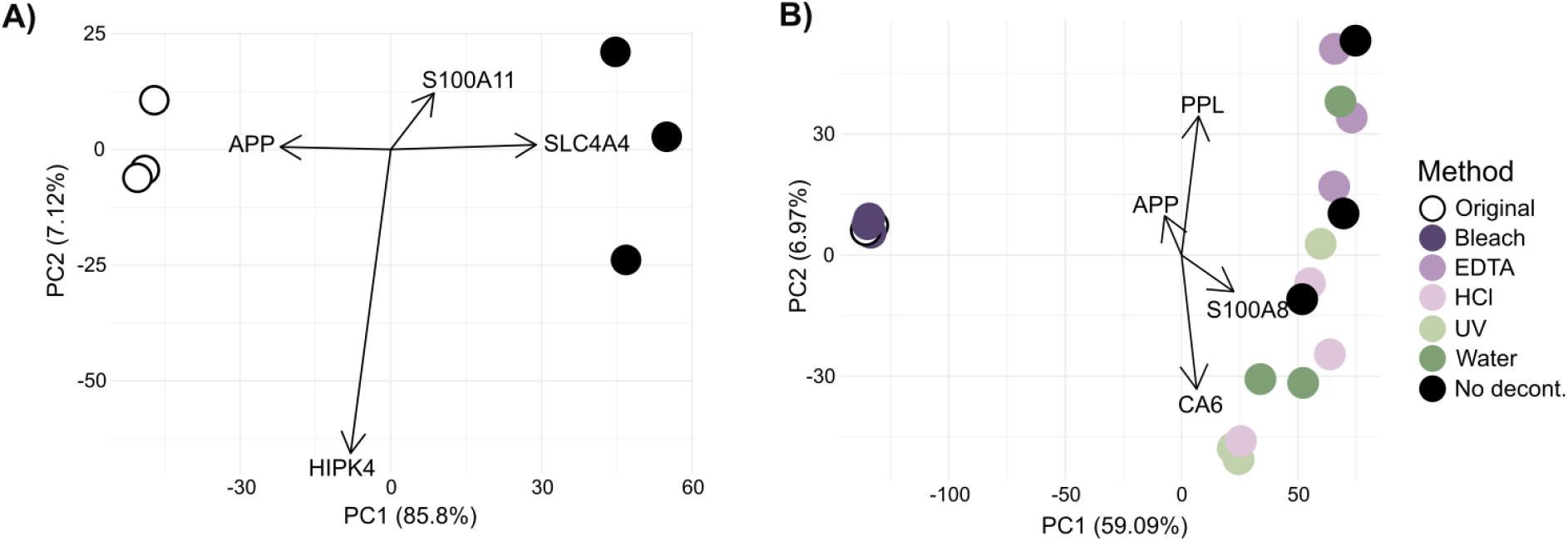
Contaminating and decontaminating a Pleistocene bone proteome. A) A PCA based on protein quantification shows separation of contaminated and non-contaminated (original) skeletal proteomes. Both conditions were searched together against a horse protein database, without entries for dog proteins present, and without subsequent filtering of identified proteins. B) A PCA of protein intensities based on a database search using both dog and horse proteomes as reference. In both PCAs, gene names of proteins contributing most to the variation along each axis are shown; S100A11=protein S100A11, APP=amyloid-beta precursor protein, SLC4A4=anion exchange protein, HIPK4=homeodomain-interacting protein kinase 4, PPL=periplakin, S100A8=protein S100A8, CA6=carbonic anhydrase 6.

The number of acquired MS2 spectra is 25,931±148 for the original samples and 23,756±203 for the contaminated samples, with significant differences between the sample groups (ANOVA, F=225.3, p<0.001). Additionally, the percentage of identified MS2 spectra differs significantly between the original and contaminated samples (ANOVA, F=9.24, p=0.038), with a higher percentage of MS2 spectra being identified in the original samples (5.10±0.46%) than the contaminated samples (4.13±0.31%). Together, this indicates that the presence of contaminants is significantly reducing the amount of endogenous spectra that can be acquired and identified. We therefore conclude that protein contamination has a relevant impact on the composition of reconstructed endogenous skeletal proteomes, even when the protein contamination is not considered or identified.

Next, we searched all extracts from the five decontamination methods, the non-decontaminated condition and the original samples against a database containing the complete dog and horse proteomes (including the MaxQuant contaminant database). For the non-decontaminated samples, 154.0±9.7 proteins and 1,481±104.9 PSMs were identified, whereas for the original sample, 43.0±3.5 proteins and 1,364±135.7 PSMs were identified. A PCA shows that the original and non-decontaminated extracts are clearly separated along PC1 (Figure 2B), and that the decontamination method significantly affects proteome composition (PERMANOVA, R2=0.72, p=0.001). PC1, accounting for 59.09% of variation in the dataset, separates bleached and original proteomes from the uncleaned extracts, with all other decontamination approaches placed close to the latter. The extracts resulting from UV irradiation, HCl wash, EDTA wash and molecular grade water wash fall on a cline together with the uncleaned extracts, with separation between decontamination approaches along PC2, accounting for 6.97% of variation in the dataset. The proteins driving separation of the original and bleached samples from all other conditions along PC1 are expressed in saliva (CA6; carbonic anhydrase 6), skin (PPL; periplakin), and are involved in inflammatory processes (S100A8).

After excluding taxonomically unspecific identifications, we observe that the dog proteome size, i.e. the number of dog proteins identified, is larger than the horse proteome size in our contaminated but uncleaned extracts (the ratio of dog:horse proteins being 3.29±0.30; Figure 3A), further demonstrating the successful contamination of the horse bone proteome with dog proteins. The ratio of dog:horse proteins is significantly dependent on the decontamination method (ANOVA, F=31.2, p<0.001), with bleaching and original samples having a significantly lower ratio of dog:horse proteins than every other condition (Tukey’s HSD, p<0.001 in each case), but not significantly differing from each other. The other decontamination approaches have variable impacts on the removal of protein contamination. These findings are mirrored when looking at the number of peptide-spectrum-matches (PSMs), where we observe that the decontamination methods also significantly affect the ratio of dog:horse PSMs (ANOVA, F=29.51, p<0.001; Figure 3B), with bleach (0.39±0.02) being similar to the original samples (0.36±0.03), and both of them significantly lower than all other methods (Tukey’s HSD, p<0.001 in each case). In the uncontaminated extracts, a small number of proteins are identified as stemming from a dog. These proteins are mainly represented by 1-2 peptides each; the ones with a higher abundance are bone proteins, and thereby likely horse proteins misidentified as stemming from a dog. The increased number of horse proteins identified in all conditions except for bleach do not represent bone proteins; the number of bone-derived proteins stay constant across all conditions. There is no significant difference in the total intensity of horse-derived proteins (normalized by weight of input material) between the decontamination methods (ANOVA, F=1.57, p=0.23).

**Figure 3.**
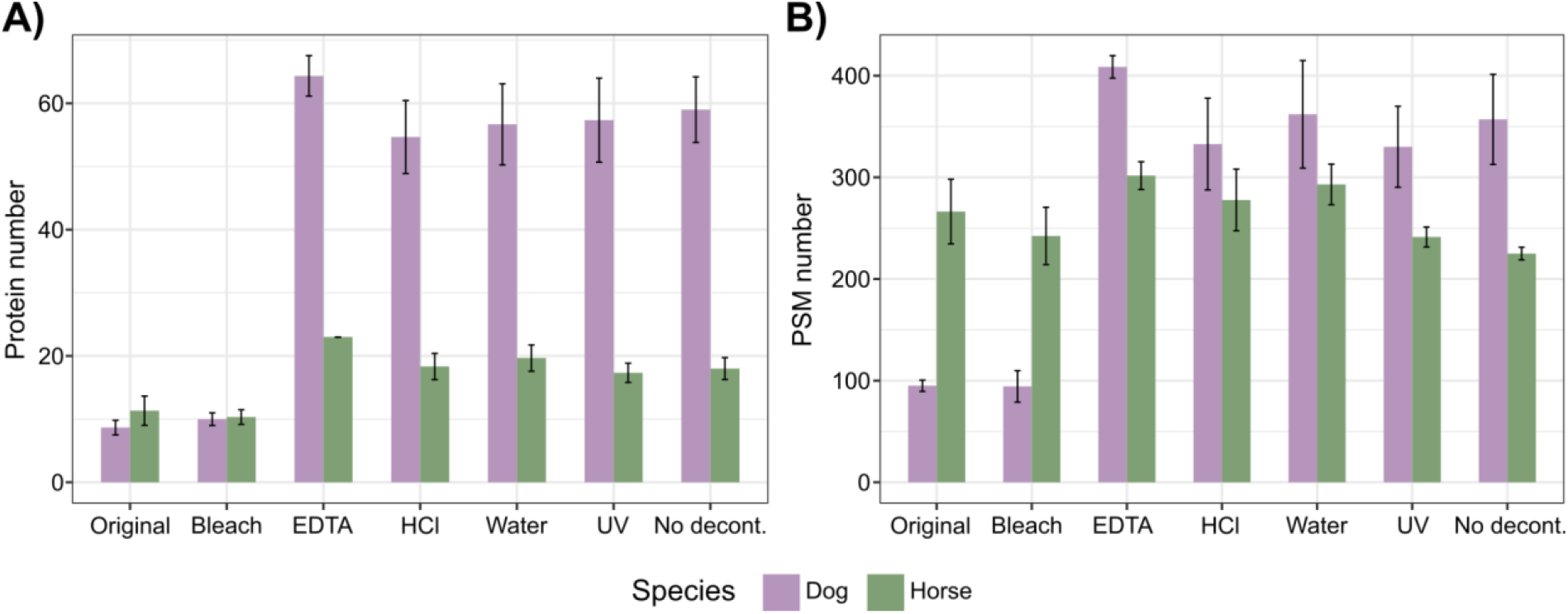
Species-specific protein recovery, in terms of A) Number of protein groups and B) Number of peptide spectral matches (PSMs). Bars show the mean of the three replicates ± 1 SD.

As an example of the effectiveness of the decontamination treatments, we examined the protein lactotransferrin (LTF, also known as lactoferrin or LF) closer. LTF is a common component of secretory fluids including saliva, and is one of most abundant dog contaminants in our dataset, but absent from the original samples. The different decontamination treatments have varying impacts on LTF abundance in terms of peptide counts or the sequence coverage obtained (Figure 4A-B). While an HCl wash has very little impact on LTF abundance, bleaching completely removes all LTF peptides from our identified dataset.

**Figure 4.**
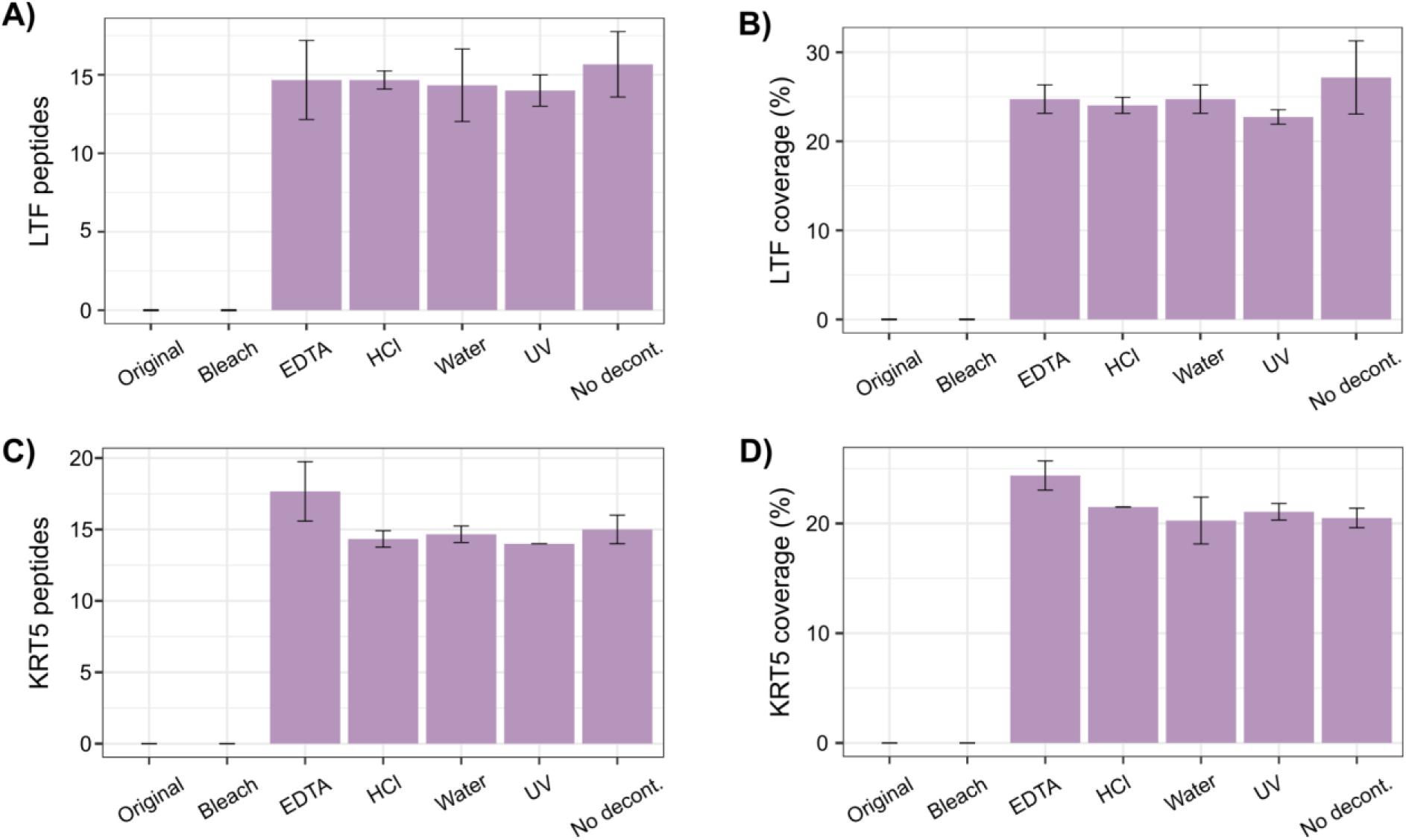
Abundance of contaminants through different decontamination methods. A) Number of dog lactotransferrin (LTF) peptides, B) Sequence coverage percentage of dog LTF, C) Number of human keratin type II cytoskeletal 5 (KRT5) peptides, and D) Sequence coverage percentage of human KRT5. Bars show a mean of the triplicate extractions, and error bars show ± 1 SD.

Since human contamination of the *Equus* bone is also possible, e.g. during excavation or handling of the specimen as well as during the dog protein contamination event, we investigated the presence of human proteins identified through the contaminant database included in the MaxQuant search. All identified human proteins are keratins. The largest number of human protein groups and peptides were identified in the EDTA-treated extracts (10.0±0.00 proteins, 37.7±1.15 peptides). None were identified in the original sample or after bleach-decontamination. The most common human protein, keratin type II cytoskeletal 5 (KRT5), is present at a maximum of 17.6±2.08 peptides and a sequence coverage of 24.4±1.33% in the EDTA treated extracts, whereas is is absent in the original and bleached extracts (Figure 4C-D). As no keratins are identified in the original samples, the human contamination likely occurred during the dog contamination, as it did not take place in a clean laboratory environment.

In addition to the removal of contaminating proteins, it is equally important to know if and how the decontamination methods affect endogenous proteins and peptides, to ensure that the ancient proteins are not damaged further. We find that coverage across a highly abundant protein (collagen alpha-2(I) chain; COL1A2) and a common non-collagenous protein (chondroadherin; CHAD) are not systematically affected by any decontamination approach (Figure 5A-B), indicating that the decontamination methods do not hinder the recovery of endogenous proteins. Some regions of COL1A2 are, however, only accessible through decontamination. The number of recovered amino acid positions of COL1A2 is affected by decontamination method (ANOVA, F=7.40, p=0.001), with significant differences detected between UV and bleach, EDTA, HCl, water and no decontamination, as well as between no decontamination and water (Tukey’s HSD, p<0.05 in each case). Overall, UV treatment leads to the lowest number of recovered amino acid positions (526±2.89) and a water wash to the highest (611±9.85). Furthermore, when comparing the non-decontaminated extracts for CHAD with the different decontamination methods, it is apparent that decontamination increases the recovery of this low-abundance protein. The highest number of recovered amino acid positions of CHAD is from the original samples (128±8.02) and the lowest through the non-decontaminated samples (92.0±1.73). The number of recovered amino acid positions depends on the method (ANOVA, F=3.47, p=0.026) with significant differences between no decontamination and the original as well as HCl-treated samples (Tukey’s HSD, p=0.039 in both cases). Lower-abundance proteins or protein regions may therefore only be accessible after decontamination.

**Figure 5.**
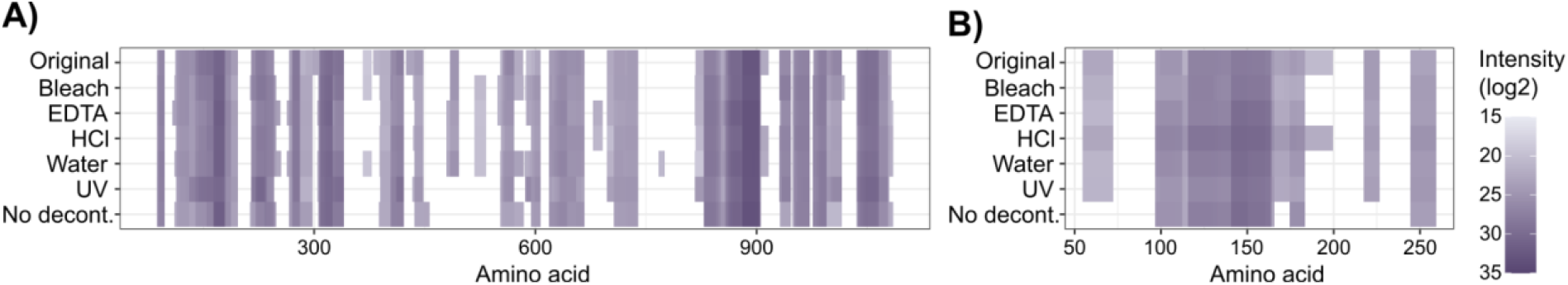
Coverage of horse bone proteins after the different decontamination methods. Coverage is shown as log2 of the summed LFQ intensity per amino acid position, averaged across the triplicate extracts, for A) COL1A2 (collagen alpha-2(I) chain) and B) CHAD (chondroadherin).

Endogenous peptide length is significantly affected by the decontamination methods (ANOVA, F=3.08, p=0.039; Figure S2A), however, the only significant difference between treatments is between UV irradiation and no decontamination (Tukey’s HSD, p=0.022), while none of the decontamination methods show significant differences in comparison to the original sample. The post-translational modifications (PTMs) that were explored, including hydroxyproline, deamidation and UV-specific modifications, are not affected by the decontamination methods tested either (ANOVA, p>0.05 for all; Figure S2B-E). It can therefore be concluded that even the most efficient decontamination methods do not damage the ancient endogenous proteome by any of the metrics examined here.

From this we conclude that all decontamination approaches tested have some positive impact on the removal of contaminating proteins, but to various extents and with bleach being the only approach that results in the recovery of a skeletal proteome highly similar to the endogenous, uncontaminated proteome of the original bone specimen.

### 3.2. Application of the most promising method to a contaminated hominin proteome

Artificial contamination of a bone fragment was used to determine the most efficient decontamination method; however, this is only an approximation of actual contamination of an archaeological object over time. During initial analysis of the dentine proteome of a Pleistocene hominin tooth from Khudji, Tajikistan (Figure 6A), we noted that the proteome was heavily contaminated with proteins originating from human skin. Bone and dentine are quite similar in terms of composition, both consisting mainly of the biomineral hydroxyapatite and having similar ratios of organic to inorganic material, although dentine has lower porosity than bone (Kendall et al., 2018). The same decontamination protocol should thus be applicable to dentine. The bleach decontamination protocol was therefore applied to a second dentine fragment from the tooth. Of the 34 protein groups recovered from the initial dentine proteome analysis, we classified six of these as endogenous dentine proteins based on the PhyloBone database (Fontcuberta-Rigo et al., 2023) (Figure 6B). Of the remaining protein groups, 21 derived from skin, and seven from other tissues (including proteins that are present in several tissues). In contrast, after bleach treatment, eight protein groups originating from dentine were recovered, and only one each from skin and other tissues. A smaller total number of peptides were recovered after bleach treatment, but the vast majority of them originate from dentine proteins. Although there is a difference in peptide numbers identified, it should be noted that the bleached and untreated extracts derive from different dental fragments, different sample weights, and have different numbers of acquired MS2 spectra. As a result, we cannot directly compare the extracts based on absolute numbers, but only describe their proteomic composition in relative terms. The pattern is, however, the same as observed for the controlled contamination-decontamination experiment outlined above, where bleach treatment significantly lowered the ratio of contaminant:endogenous proteins. In the untreated Khudji extract, the ratio of contaminant to endogenous (skin:dentine) proteins is 3.50, while in the bleached extract it is 0.13. The same pattern is present on the peptide level, with a ratio of 0.27 for the untreated extract, and 0.01 for the bleached extract. Decontamination by bleaching thereby significantly lowers the proportion of the reconstructed proteome that stems from contamination, and allows for reconstruction of an authentic ancient dentine proteome.

**Figure 6.**
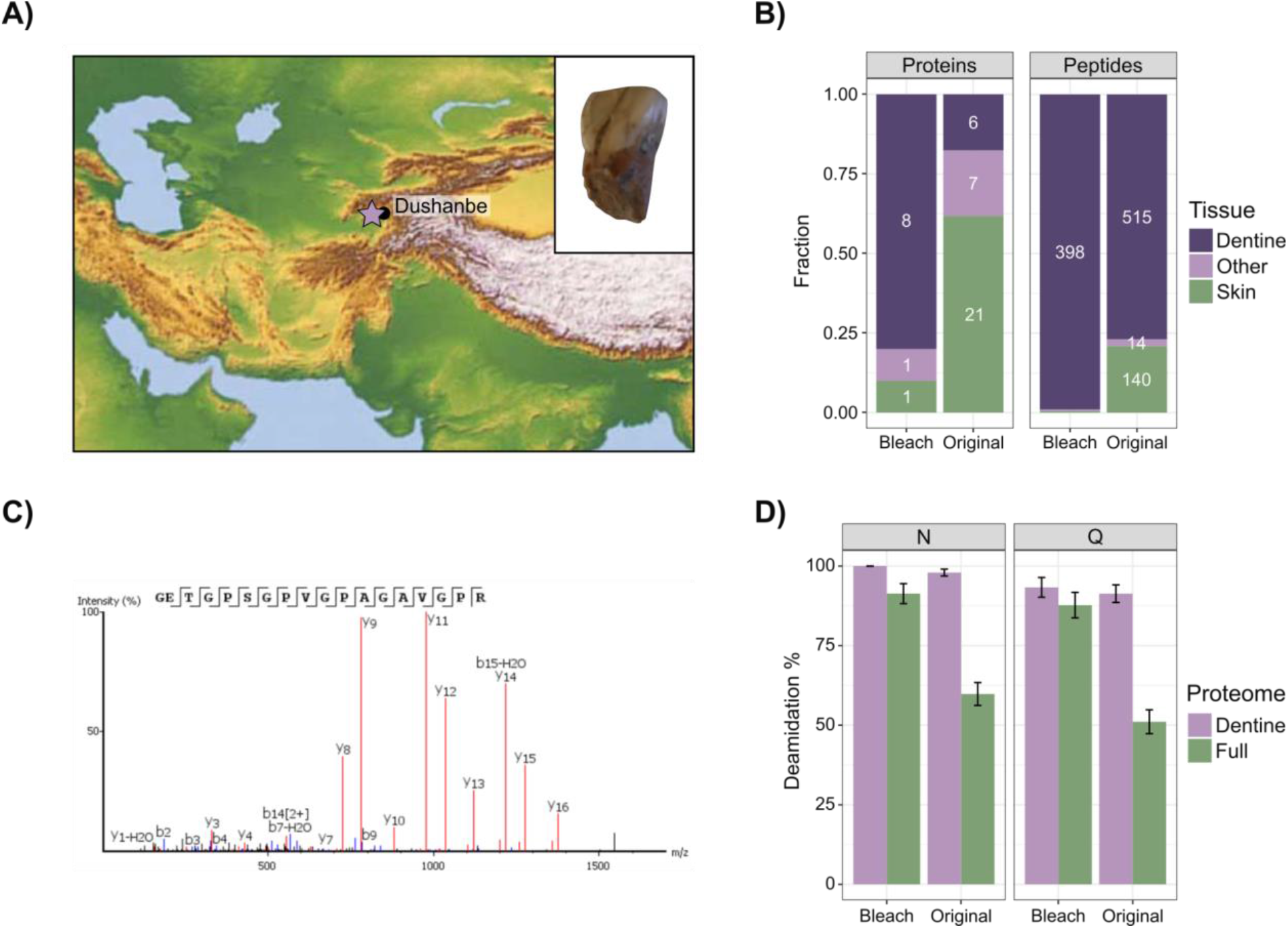
The Khudji hominin dentine proteome. A) Location of Khudji (indicated by star) and photo of the specimen. B) Proteome composition, with the fraction of proteins and peptides originating from dentine, skin, or other tissues. Numbers show counts of proteins and peptides, respectively. C) Representative MS2 spectrum covering the COL1A2 R996K position for the Khudji hominin. D) Deamidation of N and Q, with error bars showing ± 1 standard deviation. Here, 0% indicates no deamidation while 100% indicates complete deamidation of the respective amino acid.

Similar to the decontamination experiment, we quantified the extent of various protein alterations to document whether bleach treatment has a negative impact on the endogenous proteome of the Khudji dentine sample. The peptide length distribution is similar between the full bleached and unbleached proteomes (linear model, weighted by intensity; F=0.03, p=0.86; Figure S3A), but the dentine-fraction of the bleached proteome has a higher weighted mean peptide length compared to the unbleached dentine proteome (linear model, weighted by intensity; F=22.91, p<0.001). The peptides in the unbleached sample have higher hydrophobicity than in the bleached sample (linear model, weighted by intensity; F=14.07, p<0.001; Figure S3B), but hydrophobicity of the peptides does not differ between the dentine-derived protein fractions of the two treatments (linear model, weighted by intensity; F=0.72, p=0.39). Deamidation of both N and Q is higher in the bleached extract for the full proteome, indicating a higher proportion of ancient, endogenous peptides (Figure 6D). Together, this shows that also when applied to a Pleistocene human proteome, the bleach decontamination protocol is highly efficient at removing modern contaminants, without damaging the endogenous human proteome.

The Khudji tooth was previously morphologically identified as stemming from either a modern human or a Neanderthal (Trinkaus et al., 2000). The taxonomic identity of the Khudji specimen has been subject of discussion since then, with the subsequent discovery of Denisovans adding a third possibility. In order to achieve a molecular taxonomic identification, protein sequences were reconstructed from both the bleached and untreated extracts of the Khudji hominin tooth, for all proteins with more than five peptides (SI Table 1; Supplementary Data 2). For the untreated extract, this led to nine proteins being reconstructed, with a coverage of 2.2-66.5%, whereas for the bleached extract, six proteins with a coverage of 2.5-53.3% were reconstructed. For phylogenetic analysis, the two reconstructions were combined, to form a consensus protein sequence reconstruction for the Khudji individual with the maximal amount of information included. This led to the phylogenetic analysis being based on 12 proteins, with a total coverage of 3,041 identified amino acid positions. The Khudji individual does not have coverage of any Neanderthal-specific single amino acid polymorphisms (SAPs), but lacks a Denisovan-derived SAP at COL1A2 R996K (Chen et al., 2019) (the position is R modern humans and Neanderthals, and K in Denisovans). For this position, the Khudji individual retained the ancestral R (Figure 6C). We therefore determine that, among the reference protein sequences currently available, and within the constraints of the protein sequence coverage obtained, the Khudji tooth is likely not a Denisovan, but further taxonomic determination is not possible at present.

## 4. Discussion

Previous palaeoproteomic studies have either assumed that decontamination is not necessary, or they have applied a decontamination method without estimating its efficiency at removing contaminating proteins, nor assessing its impact on endogenous proteins. We show here that protein contamination can significantly impact downstream palaeoproteomic analyses, even when the contaminating proteins stem from a different tissue and a different species. The reconstructed horse skeletal proteome, which was contaminated with dog saliva and skin, has a different composition to the original skeletal proteome, even when the potential of dog proteins being present in the dataset is not considered (Figure 2A). Further, we show that endogenous, low-abundance proteins and protein regions may not be detected in contaminated samples, likely due to the higher abundance of contaminant proteins. As the impact of protein contamination is clear even between distantly related taxa, it is increasingly important to consider the impact that human contamination may have on analysis of archaeological hominin samples.

We evaluate a range of commonly used decontamination methods, both in terms of contaminant removal and their impact on the endogenous proteome. These methods were selected based on being commonly used in the palaeoproteomics field, without taking into account their theoretical efficiency in protein contaminant removal. Many of the methods do remove protein contamination to some degree, especially when focusing on protein abundance, rather than presence/absence of contaminant proteins. However, only a single one of the tested methods seemingly returns the proteome to its original state - washing with a 0.5% solution of bleach (sodium hypochlorite). Bleach is commonly used as a disinfectant, due to its ability to disrupt cells and degrade biomolecules (Ersoy et al., 2019; Fukuzaki, 2006). A brief wash with HCl or EDTA will, on the other hand, demineralize the surface layer of the bone, thereby releasing exogenous proteins bound to the mineral surface. The UV irradiation method commonly used in published palaeoproteomics literature may have been inefficient due to the short duration of the irradiation, combined with the fact that the saliva had dried and absorbed to the bone (Moore et al., 2011; Sagripanti & Lytle, 2011). Finally, the water wash likely only removes particles that are not bound to the mineral surface.

Subsequent to bleach treatment, the reconstructed proteomes closely resemble those of uncontaminated samples. However, it is also important to investigate the damage caused by decontamination protocols. A harsh decontamination method may remove all contaminating proteins, but could also damage the already fragmented and modified ancient proteins. Of the methods tested and preservation parameters considered here, none cause significant damage to the endogenous proteins. This indicates that even though a bleach wash removes surface contamination, it is not able to access the endogenous proteins in the bone.

Although the artificial contamination method chosen for this study - modern dog saliva and skin - is unconventional, it appears to have been successful. Given that the contamination is not removed by a water wash, it has absorbed well enough to the bone to mimic long-term contamination from handling. A large number of dog proteins are identified, with the most common ones being typical for saliva or keratins. Given that the protein extraction protocol employed in this study is adapted to archaeological materials, and thereby does not contain steps such as cell lysis to release proteins, the abundance of recovered dog-derived proteins may be reduced compared to what was introduced to the bone during contamination. However, as this situation mimics actual contamination and processing of an archaeological bone, through e.g. contact with human skin followed by extraction of ancient proteins, the results approximate authentic contamination and decontamination scenarios in palaeoproteomic studies.

Application of the most promising decontamination method, a brief bleach wash, to a Pleistocene hominin tooth showed that bleach treatment is also successful in a non-artificial contamination context. The extracted proteome initially consisted of a large amount of skin-derived proteins, likely stemming from handling since excavation of the specimen in 1997 (Trinkaus et al., 2000). After bleach treatment, a proteome consisting almost solely of dentine-derived proteins was reconstructed. Further, the endogenous ancient hominin proteome appeared to not have been damaged by the bleach treatment. Analysis of the reconstructed protein sequences from both extracts from this tooth showed that it likely does not stem from a Denisovan, but further taxonomic resolution cannot be obtained from the data currently available. In the geographic region around Khudji, Neanderthal fossils have been conclusively identified at Teshik-Tash (Krause et al., 2007) and Obi-Rakhmat (Bailey et al., 2008), with further possible Neanderthal fossils recovered from Sel’Ungur and Anghilak (M. Glantz et al., 2008; Krivoshapkin et al., 2020). In addition, the Mousterian lithic technology recovered from Khudji is represented at other archaeological sites in the region (M. M. Glantz, 2011; Khujageldiev & Kunitake, 2023; Krivoshapkin et al., 2020; Ranov & Amosova, 1984; Ranov & Khujageldiev, 2014). Given the complex occupational histories recovered from the southern Altai, further east, where across the Late Pleistocene the presence of Neanderthals, Denisovans, and modern humans is attested (Kuzmin et al., 2022; Skov et al., 2022; Zavala et al., 2021), a similarly complex scenario might exist for Central Asia.

Given the impact of protein contamination on downstream palaeoproteomic analysis of archaeological tissues, and the success of bleach decontamination identified in the present study, we recommend that researchers studying ancient proteins in other types of archaeological tissues conduct similar tests. As different archaeological tissues, such as enamel, dental calculus, or ceramic food crusts, have different compositions of both organic and inorganic components, they may require other types of decontamination approaches for optimal results. Finally, by applying this approach to a Pleistocene hominin tooth containing abundant human skin protein contaminants, we demonstrate that brief bleaching opens up the analysis of Pleistocene hominin specimens otherwise inaccessible to palaeoproteomic analysis.

## 5. Conclusions

The existence of modern protein contamination is a recognised issue in the field of palaeoproteomics, but to-date no comparative studies have explored the consequence of this modern protein contamination, nor tested the efficiency of published decontamination procedures in both removing protein contamination and retaining endogenous proteomic information. Firstly, we demonstrate that the presence of protein contamination has an impact on the identification of endogenous peptides. Secondly, we provide evidence that a brief bleach wash removes protein contamination without further damaging the endogenous peptides in a controlled experimental setting. Subsequently, we demonstrate that this approach can also be applied to a Pleistocene hominin tooth. Since protein contamination is a recurring but understudied aspect in palaeoproteomic studies, we believe that our approach paves the way for the study of archaeological specimens that are heavily contaminated with modern (human) proteins, for example through long-term and extensive handling in curatorial facilities.

## Data availability

Proteomic data has been deposited to the ProteomeXchange Consortium via the PRIDE (Perez-Riverol et al., 2022) partner repository with the dataset identifier PXD050393 (*Equus* sp.), and PXD050370 and 10.6019/PXD050370 (*Homo* sp.). R code used for analysis can be found at https://github.com/ZandraFagernas/bone_decontamination.

## Acknowledgements

This research has been made possible through funding from the European Research Council (ERC) under the European Union’s Horizon 2020 research and innovation programme, grant agreement no. 948365 (PROSPER, awarded to F.W.), the European Union’s Horizon Europe research and innovation programme under the Marie Skłodowska-Curie grant agreement no. 101106627 (PROMISE, awarded to Z.F.), and through generous funding from the Leakey Foundation (awarded to Z.F.). Views and opinions expressed are those of the author(s) only and do not necessarily reflect those of the European Union or the European Research Council Executive Agency. Work at the Novo Nordisk Foundation Center for Protein Research is funded in part by a donation from the Novo Nordisk Foundation (grant number NNF14CC0001). The Khudji tooth was found during excavations in 1997, funded by the US National Geographic Society (grant 5915-97). The analysis of archaeological material from the Khudji site was carried out with the help of grant 0121TJ1212 «History of the Tajik people». This work was partly supported by NordForsk through the funding to ‘The timing and ecology of the human occupation of Central Asia’, project number 105204. We also thank Pontus Skoglund and Oded Rimon for valuable feedback on the first version of this manuscript. Finally, we thank the Contaminator, Tjorven (Tastaway’s Herrmann), for his contribution to the research.

## Author contributions

Z.F. and F.W. designed research; Z.F. and G.T. performed research; J.V.O. contributed new reagents/analytic tools; J.P.B, T.K., R.K. and M.W.P. provided material; Z.F. and V.V.I. analyzed data; Z.F. and F.W. wrote the paper with input from the other co-authors.

