## Supplementary Figures and Tables for "Cleaning the Dead: Optimized decontamination enhances palaeoproteomic analyses of Pleistocene skeletal material"

AFFILIATIONS
1. Globe Institute, University of Copenhagen, Copenhagen, Denmark 1353
2. Department of Physics, Technical University of Denmark, Roskilde, Denmark 4000
3. Institute of History, Archaeology and Ethnography, National Academy of Sciences of Tajikistan, Dushanbe, Tajikistan 734025
4. Novo Nordisk Foundation Center for Protein Research, University of Copenhagen, Copenhagen, Denmark 2200

Corresponding Author:

Frido Welker
Øster Farimagsgade 5
Building 7, 1st floor
1353, Copenhagen
Denmark
+45 35 32 64 22


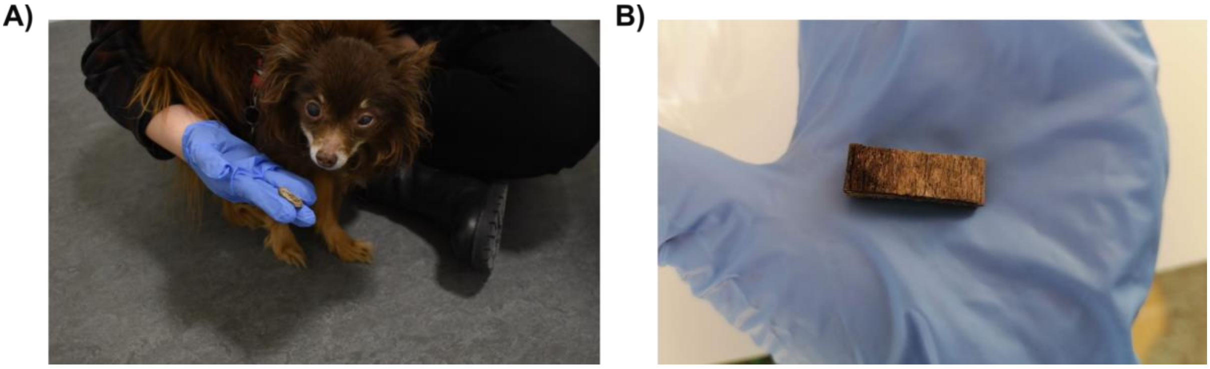


Figure S1. Contamination of a Pleistocene *Equus* sp. bone fragment. A) Contamination by saliva and fur of a dog. B) Contaminated bone fragment, where the dark end, covered in saliva, was homogenized to a single powder and subsampled for the experiment.

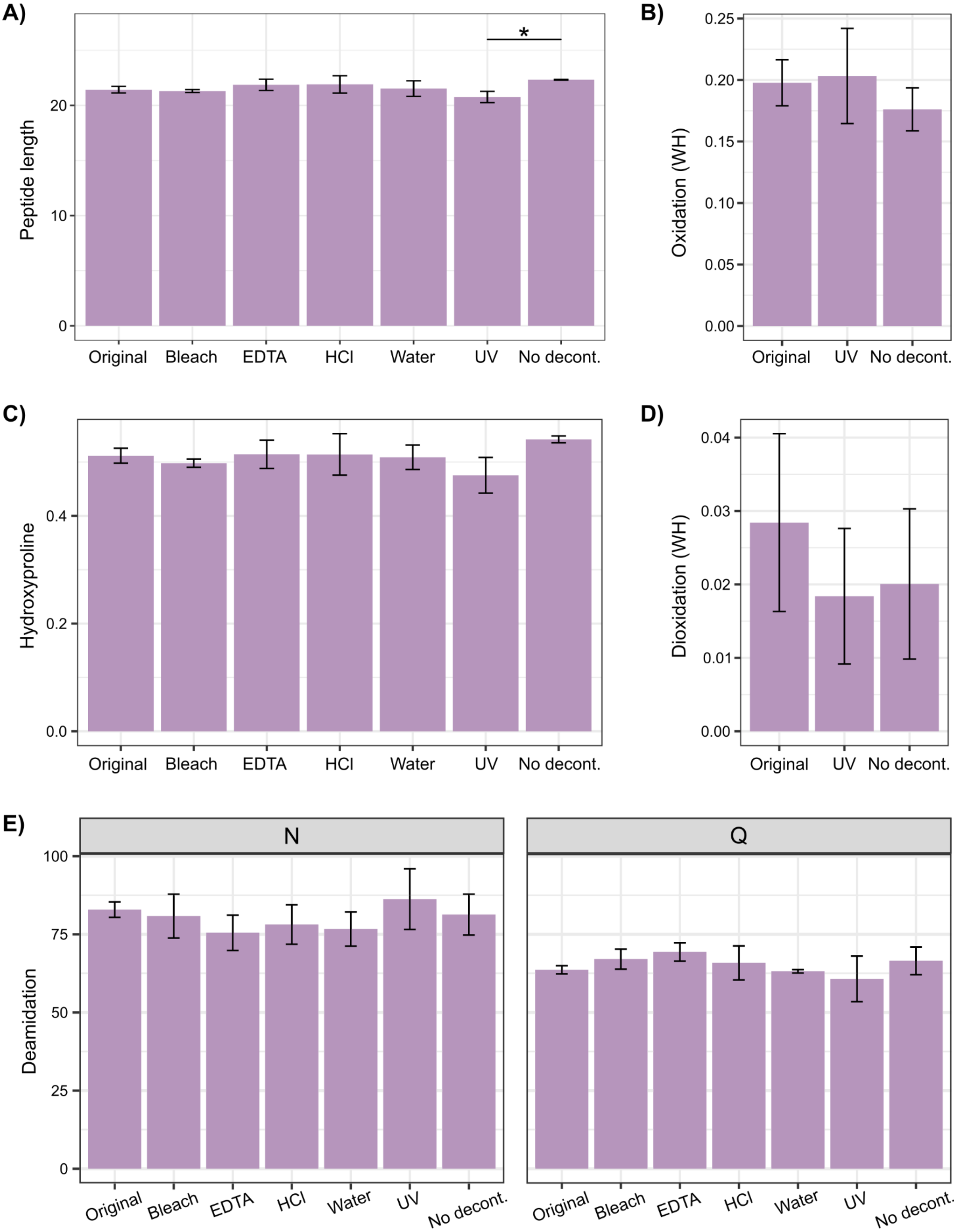


Figure S2. State of endogenous horse peptides. A) Peptide length weighted by peptide intensity. B) Oxidation of tryptophan and histidine (WH). C) Hydroxyproline. D) Dioxidation of tryptophan and histidine (WH). E) Deamidation of asparagine and glutamine (NQ). All PTMs in panels B to D are expressed as the fraction of the specific amino acids with modifications, weighted by peptide intensity. Error bars show ± 1 SD calculated from the three replicates per method. Significant differences between methods are indicated by asterisks (*p < 0.05; **p < 0.01; ***p < 0.001).

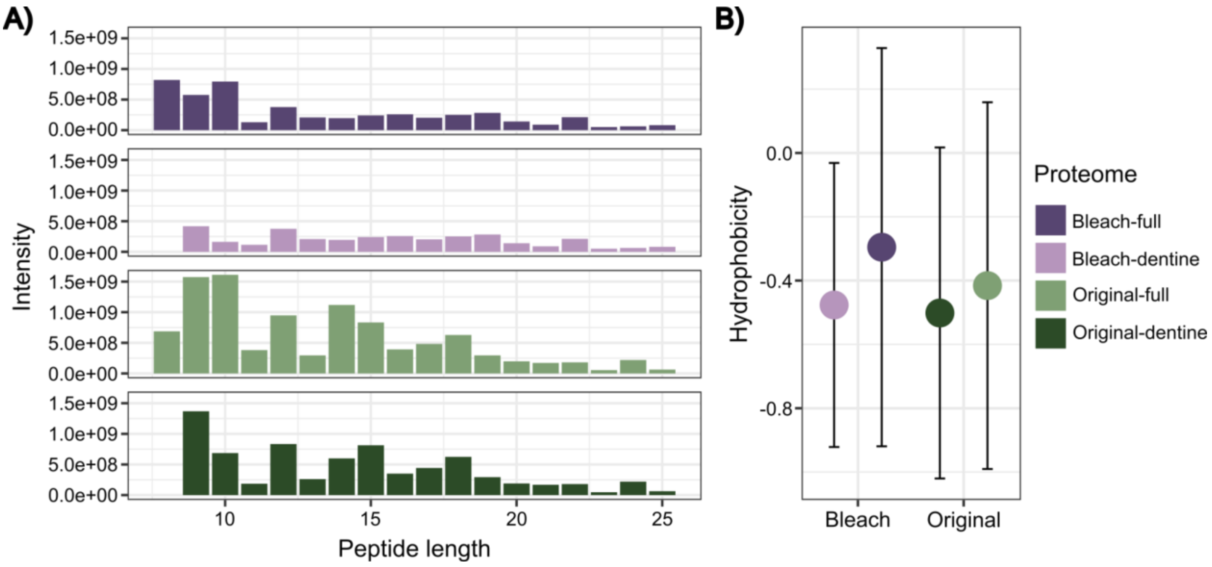


Figure S3. Properties of the Khudji dentine proteome. A) Summed intensity of each peptide length from bleached and unbleached extracts, shown both for the full proteome ("full") and when restricted to skeletal proteins only ("dentine"). B) Peptide hydrophobicity weighted by intensity. Error bars show ± 1 SD.

Table S1. Reconstructed protein sequence coverage of the Khudji dentine proteome, separated into the bleach-treated extract, the untreated extract, and a consensus sequence reconstructed from both extracts together. Percentage coverage is calculated based on the corresponding modern human protein, and the total percent coverage is calculated based on the complete concatenated sequence of proteins. It should be noted that the untreated and bleached extracts cannot be directly and quantitatively compared due to differences in the material that is analyzed (different fragments and weights) as well as different numbers of acquired MS2 spectra.

| **Protein** | **Untreated** | | **Bleach** | | **Consensus** | |
|---|---|---|---|---|---|---|
| | **Amino acids** | **Coverage** | **Amino acids** | **Coverage** | **Amino acids** | **Coverage** |
| COL1A1 | 973 | 66.5% | 781 | 53.3% | 977 | 66.7% |
| COL1A2 | 898 | 65.7% | 729 | 53.3% | 899 | 65.8% |
| COL2A1 | 327 | 22.0% | 225 | 15.1% | 385 | 25.9% |
| COL3A1 | 247 | 16.8% | - | - | 247 | 16.8% |
| COL4A1 | 37 | 2.2% | - | - | 37 | 2.2% |
| COL4A3 | - | - | 41 | 2.5% | 41 | 2.5% |
| COL4A4 | - | - | 42 | 2.5% | 42 | 2.5% |
| COL4A5 | 91 | 5.4% | - | - | 91 | 5.4% |
| COL5A2 | 158 | 10.5% | - | - | 158 | 10.5% |
| COL10A1 | 62 | 9.1% | - | - | 62 | 9.1% |
| COL18A1 | - | - | 53 | 3.0% | 53 | 3.0% |
| COL24A1 | 49 | 2.9% | - | - | 49 | 2.9% |
| Total | 2842 | 21.8% | 1871 | 19.8% | 3041 | 16.8% |
