## Supplementary Information A for "Cleaning the Dead: Optimized decontamination enhances palaeoproteomic analyses of Pleistocene skeletal material"

### A.1. The Khudji site

The Late Pleistocene site of Khudji is located in the village of Khudjii Bolo (Shakrinac district) near the city of Dushanbe, at 38°36’1’’N, 68°25’46’’E and 1,095 m above sea level (Khujageldiev & Kunitake, 2023; Ranov & Amosova, 1984; Ranov & Khujageldiev, 2014). The first finds on the surface of the slope were discovered in 1953, and excavations subsequently carried out in 1978, 1997 and 2019. The cultural layer of the site is confined to buried soils and contains four levels of archaeological material. The total number of finds at the site is about 14,000 stone artifacts and 5,000 faunal bone specimens. The Khudji lithic industry belongs to the Levallois Mousterian facies of the Middle Paleolithic of Central Asia, and is characteristic of the transitional industries from the Mousterian to the Upper Paleolithic. The main part of the artefacts consists of cores, blades, points, flakes and waste flakes. The fauna is represented by fragmented bones of caprines (*Capra/Ovis*), deer (*Alces* sp., *Cervus* sp.), horses (*Equus* sp.), tortoises (*Testudio* sp.), porcupines (*Hystrix* sp.), bears (*Ursus* sp.), bovines (*Bos* sp.), and wolves (*Canis* sp.). Palynological data indicate rather cold and humid climatic conditions at the time.

### A.2. Laboratory methods

#### *A.2.1* Equus *specimen*

Proteins were extracted from the bone powder using a standard palaeoproteomic protocol (Lanigan et al., 2020). Briefly, the samples were demineralized using 1 ml of 0.5 M EDTA at room temperature under rotation. After 24 h, the supernatant was removed by centrifuging the samples for 5 min at 12 krpm, and replaced with a fresh 1 ml of 0.5 M EDTA, and demineralization was continued for another 24 h. Buffer exchange was conducted for the EDTA supernatants using 3 kDa Amicon filter columns (Sigma-Aldrich), and replaced with an extraction buffer (2 M guanidinium hydrochloride, 10 mM Tris (2-carboxyethyl) phosphine, 20 mM 2-chloroacetamide, 100 mM Tris) bringing the volume to 300 µl. The bone pellet was washed twice using 500 µl of 50 mM Tris, and 300 µl of extraction buffer was added. Thereafter, both the pellet and supernatant fractions were incubated at 80°C for 2 h under agitation. The fractions were digested separately with 0.5 µg rLysC (Promega, V1671) for 1.5 h at 37°C at 250 rpm agitation. Thereafter the fractions were diluted to 0.6 M GuHCl using a solution of 25 mM Tris, 10% ACN and water, and trypsin digestion was carried out overnight using 2 µg trypsin (Promega, V5111) at 37°C at 250 rpm agitation. Digestion was stopped by acidifying to pH 2 using 10% trifluoroacetic acid (TFA). Peptides were cleaned on in-house made C18 StageTips using a modified version of Cappellini et al. (2019). Briefly, 150 µl of methanol was centrifuged through the Stagetips, followed by 150 µl of 80% ACN, 0.1% TFA, 20% water, and finally 150 ul of 0.1% TFA in water. Thereafter peptides were centrifuged through, binding both pellet and supernatant fractions to the same StageTip, and washed twice with 150 µl of 0.1% TFA in water. StageTips were frozen at -20°C until LC-MS/MS analysis.

#### *A.2.2* Homo *specimen*

For the initial extraction, proteins were extracted from 4.5 mg of dentine using an in-solution digestion protocol adapted for highly degraded samples (Jensen et al., 2023). Briefly, the sample was demineralized overnight at 4°C using 0.5 ml of 0.5 M EDTA. Digestion was thereafter conducted by adding 0.8 µg trypsin (Promega, V5111) and incubated at 37°C under agitation overnight. The peptides were cleaned using the StageTip protocol outlined above, without acidification. A second subsample of 5.4 mg of dentine was extracted using the same protocol, with the exception of adding a bleaching step prior to demineralization, as described above for the *Equus* sp. sample, and demineralization at room temperature under rotation.

### A.3. Mass spectrometry

A 33 µl solution of 40% ACN, 0.1% TFA was used to elute peptides from StageTips, whereafter the extracts were evaporated to approximately 1 µl, and reconstituted in 20 µl of 0.1% TFA, 5% ACN. An EASYnLC 1200 system was used for liquid chromatography (Thermo Fisher Scientific, Waltham, MA, USA), using mobile phases A (0.1% formic acid in water) and B (80% acetonitrile, 20% water, 0.1% formic acid). The peptide eluate was loaded at 500 bar pressure, whereafter a gradient elution was performed with two linear segments, transitioning from 5-30% solution B over 50 min, followed by a 10 min step to 45% of solution B. A wash was conducted at 80% of solution B followed by re-equilibration at 5% of solution B. A flow rate of 250 nl/min was used for separation at 40°C. The chromatographic column, housed in a silica tube (25 cm length and 0.75 mm inner diameter, with integrated ESI tip), was packed with Reprosil C18 beads with 1.9 µm diameter and 120Å pores (Dr. Maisch, Germany). Mass spectrometric analysis was conducted using an Exploris 480 (Thermo Fisher Scientific, Waltham, MA, USA) at 2 kV positive ionization mode at 275°C. MS1 scan was at an orbitrap resolution of 120,000 and 350-1400 m/z, and the MS2 scan was at 60,000 resolution, with m/z ranging from 100 to precursor m/z +20, in an argon environment at 1E-5 bar pressure with 60V collision energy. Fragmentation was conducted of the top 10 ions with intensity above 2e4 and a charge of 2-6, excluding isotopologues, with fragment ion exclusion for 20 s.

#### A.4. Data analysis

The *Equus* sp. LC-MS/MS data was analyzed using MaxQuant v.2.1.3.0 (Cox & Mann, 2008) with the dog (*Canis lupus familiaris*) reference proteome (UP000805418; downloaded on 2022-06-10) and/or horse (*Equus caballus*) reference proteome (UP000002281; downloaded on 2022-06-10). The internal MaxQuant contaminant database was also included, as it contains a range of common contaminants, including human keratins. A semi-specific search was conducted, with Oxidation (M), Deamidation (NQ), Gln/Glu->pyro-Glu, Carbamidomethylation and Hydroxyproline as variable modifications. To investigate UV-specific modifications, samples from the UV-treatment, original, and not decontaminated samples were analyzed with MaxQuant against a database consisting only of horse proteins identified in the previous analysis. This search was semi-specific, with Oxidation (M), Deamidation (NQ), Carbamidomethylation and Hydroxyproline as variable modifications, and adding Oxidation (WH) and Dioxidation (WH) as variable modifications as they have been identified as the most common UV-related modifications (Mackie et al., 2018). The search was conducted on a reduced database, as the number of modifications that were included increased the search space considerably. The *Homo* sp. data was analyzed using MaxQuant with a semi-specific search and Oxidation (M), Deamidation (NQ), Gln/Glu->pyro-Glu and Hydroxyproline as variable modifications, with the human reference proteome (UP000005640, downloaded on 2022-01-17) as database.

The output from MaxQuant was analyzed and visualized in R v.4.3.0 (R Core Team, 2023) using packages *tidyverse* v.2.0.0 (Wickham et al., 2019), *janitor* v.2.2.0 (Firke, 2023), *ggpubr* v.0.6.0 (Kassambara, 2023), *car* v.3.1.2 (Fox & Weisberg, 2019), *vegan* v.2.6.4 (Oksanen et al., 2022), *BioStrings* v.2.61.1 (Pagès et al., 2023), *ape* v.5.7.1 (Paradis & Schliep, 2019), *Peptides* v.2.4.6 (Osorio, 2015), *Hmisc* v.5.1.1 (Harrell, 2023), *MASS* v.7.3.58.4 (Venables & Ripley, 2002) and *MetBrewer* v.0.2.0 (Mills, 2022). Reverse hits were removed prior to analysis. When assigning proteins or peptide spectral matches (PSMs) to either dog or horse, matches to both species, where taxonomy cannot be ascertained, were not counted, as they are not informative for this study. Statistical analyses were conducted using one-way ANOVAs, and homogeneity of variances was tested using Levene’s test. Normality of residuals of the ANOVA was tested using a Shapiro test. Significant differences between pairs of methods were identified using Tukey’s HSD post-hoc tests. For PSM counts, the evidence.txt file was used, and for protein counts, the proteinGroups.txt file. Intensity-based calculations are derived from LFQ-intensity. Deamidation calculation was conducted following Mackie *et al.* (2018), and the levels of other variable modifications were calculated through taking a weighted mean (by peptide intensity) of the fraction of modified amino acids by peptide. As oxidation of methionine and carbamidomethylation calculation was based on <25 peptides per sample, the results are highly variable and therefore not included here. Peptide hydrophobicity was calculated using the Kyte-Doolittle scale.

Proteome composition was compared using a principal component analysis (PCA) based on protein group intensities. Only protein groups identified in a minimum of two extracts were included, to reduce the effects of random identifications. The intensity values were log2-transformed after pseudocount zero replacement. The proteins mainly contributing to variation in positive and negative direction on each principal component are shown in the figures. Statistical testing of the drivers of variation in the PCA was conducted using a PERMANOVA based on euclidean variation and 999 permutations.

To reconstruct the proteome from the Khudji *Homo* sp. tooth, the raw data was analyzed using PEAKS v.7.0 (Zhang et al., 2012) with a database consisting of the human reference proteome (UP000005640, downloaded on 2022-02-22) and added archaic variation from Neanderthals (Castellano et al., 2014) and a Denisovan (Meyer et al., 2012). Parent mass error tolerance was set to 10.0 ppm and fragment mass error tolerance to 0.07 Da. Variable modifications were deamidation (NQ), hydroxylation (P) and oxidation (M). Protein sequences were reconstructed separately for the original and bleached extraction, for proteins with more than five peptides, based on the PEAKS search (filtered for 0.5% FDR, following published recommendations (Welker, 2018)). A majority consensus was called at each amino acid position, and the sequences were aligned using Geneious Prime v.2023.2.1 with corresponding protein sequences from modern humans, Neanderthals (Castellano et al., 2014) and a Denisovan (Meyer et al., 2012); any identified SAPs were manually verified in PEAKS. A consensus sequence for the individual was then created (Supplementary Data 2).

### A.5. References

Cappellini, E., Welker, F., Pandolfi, L., Ramos-Madrigal, J., Samodova, D., Rüther, P. L., Fotakis, A. K., Lyon, D., Moreno-Mayar, J. V., Bukhsianidze, M., Rakownikow Jersie-Christensen, R., Mackie, M., Ginolhac, A., Ferring, R., Tappen, M., Palkopoulou, E., Dickinson, M. R., Stafford, T. W., Jr, Chan, Y. L., … Willerslev, E. (2019). Early Pleistocene enamel proteome from Dmanisi resolves *Stephanorhinus* phylogeny. *Nature*, 574(7776), 103–107.

Castellano, S., Parra, G., Sánchez-Quinto, F. A., Racimo, F., Kuhlwilm, M., Kircher, M., Sawyer, S., Fu, Q., Heinze, A., Nickel, B., Dabney, J., Siebauer, M., White, L., Burbano, H. A., Renaud, G., Stenzel, U., Lalueza-Fox, C., de la Rasilla, M., Rosas, A., … Pääbo, S. (2014). Patterns of coding variation in the complete exomes of three Neandertals. *Proceedings of the National Academy of Sciences of the United States of America*, 111(18), 6666–6671.

Chen, F., Welker, F., Shen, C.-C., Bailey, S. E., Bergmann, I., Davis, S., Xia, H., Wang, H., Fischer, R., Freidline, S. E., Yu, T.-L., Skinner, M. M., Stelzer, S., Dong, G., Fu, Q., Dong, G., Wang, J., Zhang, D., & Hublin, J.-J. (2019). A late Middle Pleistocene Denisovan mandible from the Tibetan Plateau. *Nature*, 569(7756), 409–412.

Cox, J., & Mann, M. (2008). MaxQuant enables high peptide identification rates, individualized p.p.b.-range mass accuracies and proteome-wide protein quantification. *Nature Biotechnology*, 26(12), 1367–1372.

Firke, S. (2023). janitor: Simple Tools for Examining and Cleaning Dirty Data.

Fox, J., & Weisberg, S. (2019). An R Companion to Applied Regression, Third Edition. Sage.

Harrell, F. E., Jr. (2023). Hmisc: Harrell Miscellaneous.

Jensen, T. Z. T., Yeomans, L., Le Meillour, L., Wistoft Nielsen, P., Ramsøe, M., Mackie, M., Bangsgaard, P., Kinzel, M., Thuesen, I., Collins, M. J., & Taurozzi, A. J. (2023). Tryps-IN: A streamlined palaeoproteomics workflow enables ZooMS analysis of 10,000-year-old petrous bones from Jordan rift-valley. *Journal of Archaeological Science: Reports*, 52, 104238.

Kassambara, A. (2023). ggpubr: “ggplot2” Based Publication Ready Plots.

Khujageldiev, T. U., & Kunitake, S. (2023). The excavation of Mousterian site Khudji in 2019 (in Russian). *Archaeological Work in Tajikistan*, 43, 13–40.

Lanigan, L. T., Mackie, M., Feine, S., Hublin, J.-J., Schmitz, R. W., Wilcke, A., Collins, M. J.,

Cappellini, E., Olsen, J. V., Taurozzi, A. J., & Welker, F. (2020). Multi-protease analysis of Pleistocene bone proteomes. *Journal of Proteomics*, 228, 103889.

Mackie, M., Rüther, P., Samodova, D., Di Gianvincenzo, F., Granzotto, C., Lyon, D., Peggie, D. A., Howard, H., Harrison, L., Jensen, L. J., Olsen, J. V., & Cappellini, E. (2018). Palaeoproteomic Profiling of Conservation Layers on a 14th Century Italian Wall Painting. *Angewandte Chemie*, 57(25), 7369–7374.

Meyer, M., Kircher, M., Gansauge, M.-T., Li, H., Racimo, F., Mallick, S., Schraiber, J. G., Jay, F., Prüfer, K., de Filippo, C., Sudmant, P. H., Alkan, C., Fu, Q., Do, R., Rohland, N., Tandon, A., Siebauer, M., Green, R. E., Bryc, K., … Pääbo, S. (2012). A high-coverage genome sequence from an archaic Denisovan individual. *Science*, 338(6104), 222–226.

Mills, B. R. (2022). MetBrewer: Color Palettes Inspired by Works at the Metropolitan Museum of Art.

Oksanen, J., Simpson, G. L., Blanchet, F. G., Kindt, R., Legendre, P., Minchin, P. R., O'Hara, R. B., Solymos, P., Stevens, M. H. H., Szoecs, E., Wagner, H., Barbour, M., Bedward, M., Bolker, B., Borcard, D., Carvalho, G., Chirico, M., De Caceres, M., Durand, S., … Weedon, J. (2022). vegan: Community Ecology Package.

Osorio, D. (2015). Peptides: A package for data mining of antimicrobial peptides. *The R Journal*.

Pagès, H., Aboyoun, P., Gentleman, R., & DebRoy, S. (2023). Biostrings: Efficient manipulation of biological strings.

Paradis, E., & Schliep, K. (2019). ape 5.0: an environment for modern phylogenetics and evolutionary analyses in R. *Bioinformatics*, 35, 526–528.

Ranov, V. A., & Amosova, A. G. (1984). The excavation of Mousterian site Khudji in 1978 (in Russian). *Archaeological Work in Tajikistan*, 18, 11–58.

Ranov, V. A., & Khujageldiev, T. U. (2014). The excavation of Middle Paleolithic site Khudji in 1997 (in Russian). *Archaeological Work in Tajikistan*, 37, 273–334.

R Core Team. (2023). R: A Language and Environment for Statistical Computing. R Foundation for Statistical Computing.

Ronquist, F., Teslenko, M., van der Mark, P., Ayres, D. L., Darling, A., Höhna, S., Larget, B., Liu, L., Suchard, M. A., & Huelsenbeck, J. P. (2012). MrBayes 3.2: Efficient Bayesian Phylogenetic Inference and Model Choice Across a Large Model Space. *Systematic Biology*, 61(3), 539–542.

Stamatakis, A. (2006). RAxML-VI-HPC: maximum likelihood-based phylogenetic analyses with thousands of taxa and mixed models. *Bioinformatics* , 22(21), 2688–2690.

Venables, W. N., & Ripley, B. D. (2002). Modern Applied Statistics with S (Fourth). Springer.

Welker, F. (2018). Elucidation of cross-species proteomic effects in human and hominin bone proteome identification through a bioinformatics experiment. *BMC Evolutionary Biology*, 18(1), 23.

Wickham, H., Averick, M., Bryan, J., Chang, W., McGowan, L. D., François, R., Grolemund, G., Hayes, A., Henry, L., Hester, J., Kuhn, M., Pedersen, T. L., Miller, E., Bache, S. M., Müller, K., Ooms, J., Robinson, D., Seidel, D. P., Spinu, V., … Yutani, H. (2019). Welcome to the tidyverse. *Journal of Open Source Software*, 4(43) 1686.

Zhang, J., Xin, L., Shan, B., Chen, W., Xie, M., Yuen, D., Zhang, W., Zhang, Z., Lajoie, G. A., & Ma, B. (2012). PEAKS DB: de novo sequencing assisted database search for sensitive and accurate peptide identification. *Molecular & Cellular Proteomics: MCP*, 11(4), M111.010587.
